# Progression of chronic kidney disease in African Americans with type 2 diabetes mellitus using topology learning in electronic medical records

**DOI:** 10.1101/361956

**Authors:** Li Wang, Xufei Zheng, Lynn S. Huang, Jianzhao Xu, Fang-Chi Hsu, Shyh-Huei Chen, Maggie C.Y. Ng, Donald W. Bowden, Barry I. Freedman, Jing Su

## Abstract

**Background:** Chronic kidney disease (CKD) is a common, complex, and heterogeneous disease impacting aging populations. Determining the landscape of disease progression trajectories from midlife to senior age in a “real-world” context allows us to better understand the progression of CKD, the heterogeneity of progression patterns among the risk population, and the interactions with other clinical conditions. Genetics also plays an important role. In previous work, we and others have demonstrated that African Americans with high-risk *APOL1* genotypes are more likely to develop CKD, tend to develop CKD earlier, and the disease progresses faster. Diabetes, which is more common in African Americans, also significantly increases risk for CKD.

**Data and Method:** Electronic medical records (EMRs) were used to outline the first CKD progression trajectory roadmap for an African American population with type 2 diabetes. By linking participants in 5 genome-wide association study (GWAS) to their clinical records at Wake Forest Baptist Medical Center (WFBMC), an EMR-GWAS cohort was established (n = 1,581). Patients’ health status was described by 18 Essential Clinical Indices across 84,009 clinical encounters. A novel graph learning algorithm, Discriminative Dimensionality Reduction Tree (DDRTree) was implemented, to establish the trajectories of declines in health. Moreover, a prediction model for new patients was proposed along the learned graph structure. We annotated these trajectories with clinical and genomic features including kidney function, other major risk indices of CKD, *APOL1* genotypes, and age. The prediction power of the learned disease progression trajectories was further examined using the k-nearest neighbor model.

**Results:** The CKD progression trajectory roadmap revealed diverse kidney failure pathways associated with different clinical conditions. Specifically, we identified one high-risk trajectory and two low-risk trajectories. Switching pathways from low-risk trajectories to the high-risk one was associated with accelerated decline in kidney function. On this roadmap, patients with *APOL1* high-risk genotypes were enriched in the high-risk trajectory, suggesting fundamentally different disease progression mechanisms from those without *APOL1* risk genotypes. The k-nearest neighbor-based prediction showed effective prediction rate of 87%.

**Conclusion:** The CKD progression trajectory roadmap revealed novel diverse renal failure pathways in African Americans with type 2 diabetes mellitus and highlights disease progression patterns that associate with *APOL1* renal-risk genotypes.

## Introduction

As a chronic, complex, and heterogeneous disease, chronic kidney disease (CKD) challenges traditional clinical research paradigms such as those in clinical trials or observational studies. CKD is common, irreversible, and affects organs beyond the kidney. In 2011 – 2014, about 14.8% of US adults had CKD and one third of the population over age 60 had moderate to severe CKD (Stages 3-5) (*1*). The life expectancy of patients with CKD is shorter than in those with normal kidney function. For adults age 50 with moderate or more severe CKD (Stages 3-5), life expectancy is reduced by 11 years (31.3%) compared with peers lacking CKD.

A major analytic challenge is heterogeneity in populations with CKD. CKD populations demonstrate dramatically diverse progression patterns, prognosis, risk factors, and responses to clinical interventions. Genetic and ancestry-based risk factors may define particular subpopulations exhibiting unique disease patterns and interactions with other CKD comorbidities. For example, it was discovered that in the *APOL1* gene (apolipoprotein L1), the “G1” (rs73885319, S342G and rs60910145, I384M) and “G2” (rs71785313 6-bp deletion) variants (*2*-*4*) strongly associate with increased risk for non-diabetic CKD (*5*, *6*). The *APOL1* risk variants are ancestry-specific. These risk alleles are present in >50% of African Americans but rare in other ethnic groups (*3*). Other known CKD sub-populations are patients with the history of acute kidney injury (AKI) (*7*-*10*), diabetes mellitus (DM), hypertension (HTN), or cardiovascular disease (CVD). Understanding the nature of these intrinsic CKD subtypes is crucial for supporting clinical decisions such as identifying patient subtypes and determining optimal management strategies.

Another challenge is due to the complexity of CKD, which requires comprehensive clinical indices to study its risk of progression, interactions with its comorbidities, and the impact on patient health. CKD is strongly associated with presence of other chronic complex diseases: about 40% of patients with DM, 32% with HTN, and 43% with CVD have moderate (Stage 3) or worse CKD (*1*). The known risk factors for CKD progression also result from genetic, demographic, and socioeconomic status, to associated disorders such as metabolic syndrome and AKI, to medication use and treatment history.

Electronic medical records (EMRs) provide unique and valuable opportunities to address these challenges in clinical decision support for managing the risks of CKD. EMRs cover very large cohorts over decades from a wide range of aspects including demographic, clinical, financial, and socioeconomic features. Modern advances in big data science significantly improve data interoperability, making meta-analysis in cross regional or national EMR-networks possible. Thus, an EMR-based clinical decision support system has the potential to revolutionize the care of patients with CKD.

However, the challenges using EMR-related big data have impeded longitudinal studies of CKD progression. Although the EMR provides rich and comprehensive information about the population across a wide age span, for a specific clinical visit or for an individual patient, EMR data are often sparse with irregularly spaced intervals. These factors challenge traditional epidemiological approaches. The recent advance of topological learning on big data, especially a novel graph learning algorithm, Discriminative Dimensionality Reduction Tree (DDRTree) (*11*-*14*), allows building the trajectory tree according to the similarities among data points from highly scattered data. Such an approach can be used to outline the common disease progression trajectories of a population using the EMR data as a whole. This strategy can unleash the power of EMR data and avoid the limitation of traditional approaches on data completeness and regularity.

In this manuscript, we focused on the high-risk African American population with type 2 diabetes mellitus as the study cohort, linked participants from a series of genomics studies with their EMRs at Wake Forest Baptist Medical Center (WFBMC), using EMR data in 1,581 patients, included 18 essential clinical indices, 84,009 clinical encounters and over 354,398 clinical records to learn and annotate the declines in chronic health condition trajectories during aging. We examined the chronic renal function declines on these trajectories and highlighted those closely associated with patients who have *APOL1* renal-risk genotypes. We further examined the predictive power of the learned disease progression trajectories. These results cast new light on understanding the diversity of CKD progression paths and their associated health conditions.

## Data and Methods

### The WFBMC African American genotyping cohort

The cohorts from a series of studies were merged and participants linked to the WFBMC EMR according to identifiable information including name and date of birth. The genotyping cohort included 9,656 African American participants recruited from the following studies between 1993 and 2014: African American-Diabetes Heart Study (AA-DHS (*15*)), Diabetes Heart Study (DHS(*15*)), African American Type 2 Diabetes Mellitus Cohort (DM2 (*16*), African Americans with end-stage renal disease (ESRD (*17*)), Family Investigation of Nephropathy and Diabetes (FIND (*18*)), as well as additional cases and controls recruited from WFBMC during this period. In all, 4,325 participants were identified as WFBMC patients, including 524 from AA-DHS; 200 from DHS; 169 from DM2; 249 from ESRD; 98 from FIND, and an additional 1,289 WFBMC cases and 1,247 WFBMC controls from related projects.

### The WFBMC EMR cohort

By February 26, 2016, this cohort was composed of 1,646,059 individuals at the WFBMC over the past three decades. It included 14.6% who were over the age of 65, and 22.3% were African American. This patient cohort represents both local and regional populations. WFBMC is the only academic medical center within the 12-county Piedmont Triad region of northwestern North Carolina. The referral region encompasses a population of approximately 8,000,000 in North Carolina, Eastern Tennessee, South Carolina, Virginia and West Virginia. The Appalachian population, unique in its demographics, culture, and socioeconomic characteristics, is significantly represented. Therefore, WFBMC EMRs have unique scientific value for studying diverse populations with special regional characteristics.

The original WFBMC EMRs are composed of the pre-Epic “Legacy” EMRs (1985 – 2012) providing invaluable longitudinal records for chronic disease research and the Epic-based “WakeOne” EMRs (since 2012) providing modern and fast-growing EMRs for clinical research as well as the major implementation platform for delivering the eCDS4CKD clinical decision support tools.

The WFUHS Translational Data Warehouse (I2B2-based TDW) integrates both Legacy and WakeOne EMRs, implement a wide range of biomedical ontologies and terminologies for EMR presentation and annotation, and curate a high-quality data warehouse for biomedical and clinical research. The WFUHS Center of the Scalable Collaborative Infrastructure for a Learning Health System (SCILHS), the National Patient-Centered Clinical Research Network, an innovative initiative of the Patient-Centered Outcomes Research Institute (PCORI), has established a comprehensive and cutting-edge EMR system according to Unified Medical Language Systems (UMSL) as prescribed in the PCORNET Common Data Model (CDM) (www.pcornet.org/resource-center/pcornet-common-data-model) to support large scale clinical informatics and phenome mining. The WFUHS TDW (56) I2b2 installation contains over 1.4 billion facts of demographics, vital status, diagnoses, procedures, labs, and medications detailing 1.65 million patients across 30.7 million encounters.

### The Working cohort (n = 1,581)

This cohort was established by recruiting patients from the WFBMC African American genotyping cohort and the WFBMC EMR according to the following criteria:

- African American
- Available genotypic data
- Type 2 diabetes mellitus (T2D)-affected
- >= 3 EMR-derived eGFR records

### CLINIC CDM

We defined a minimum information CDM, the *<u>C</u>ommon c<u>LIN</u>ic <u>I</u>ndex for <u>C</u>hronic diseases* (CLINIC) CDM, to comprehensively outline the general health conditions during the development and progression of common chronic diseases. The CLINIC CDM is composed of the following categories:

- Demographics (3 features): date of birth (DoB), sex, self-reported race;
- Vitals (5 features): diastolic and systolic blood pressure, height, weight, and body mass index (BMI);
- Laboratory tests (14 features): alanine aminotransferase (a.k.a. serum glutamic pyruvic transaminase), aspartate aminotransferase (a.k.a. serum glutamic oxaloacetic transaminase), alkaline phosphatase (ALK), total cholesterol, low density lipoprotein cholesterol (LDL), and high density lipoprotein cholesterol (HDL), creatine kinase, estimated glomerular filtrating rate (eGFR), hemoglobin, hemoglobin A1c (HbA1c), triglycerides, international normalized ratio of prothrombin time (INR), serum creatinine, total bilirubin (TBIL), and troponin;
- Diagnosis: ICD9 and ICD10 codes;
- Procedures: HCPCS, ICD9-CM, and ICD10-PCS codes;
- Medications: RxNorm codes.

The BMI and eGFR are derived from height/weight/age and serum creatinine/age/sex/race, respectively. The CLINIC CDM is compatible with Carolinas Collaboratives Common Data Model (CDM) and PCORI CDM. The data dictionary of the CLINIC CDM is provided in Supplement S1.

### The Essential Clinical Indices of clinical encounters

We defined an 18-feature Essential Clinical Indices (Supplement Table S2) for each clinical encounter to quantitatively describe the overall health conditions of patients. These indices are composed of all vital and laboratory variables in the CLINIC CDM except BMI since BMI was highly correlated with height and weight. The selection of these clinical indices was a tradeoff of the following considerations:

- Reflect health status and risks of chronic disease with aging.
- Clinically measured, providing strong evidence.
- Generally available in EMR.
- Readiness to use. Most of these indices had already been cleaned and aggregated by CTSI at WFBH as part of the Carolinas Collaborative infrastructure.

### EMR data aggregation

The WFBMC EMR were extracted from both the WakeOne (WFBMC’s implementation of the Epic system) and pre-Epic systems. Therefore, a clinical feature is often referred by different names in the EMR. The mapping between original EMR features and the CLINIC CDM is listed in Supplement Table S3.

### Data cleaning and imputation

At record level, extreme numerical records were discarded according to the false discovery rates (FDRs) of z-test (FDR ≤ 0.01). Weight records above 500lb were also discarded. At encounter level, missing features in Essential Clinical Indices were imputed by smoothing the continuous testing values along the increase of ages before the interpolation of neighborhood values in terms of ages is used as the imputed value. After data cleaning and imputation, a learning cohort of 435 patients with 49,357 clinical encounters were established. Details can be found in Supplement Methods (Section: Imputation).

### Data availability patterns

The concordance of feature occurrence across all encounters in the Learning cohort was measured in pairwise fashion with the L-1 distance (Manhattan distance). Essential Clinical Indices were then hierarchically clustered by complete linkage method (*19*).

### Learning disease progression trajectories

To reveal the common trajectories of disease progression from a dataset consisting of many patients with multiple types of diseases, we treated all (imputed) values of the Essential Clinical Indices (ECI) from one patient at each encounter as individual data points. Our goal was to learn the relationship between these clinical encounters to reveal topological structure correlated to clustering structure or embedded disease develop structures. We used DDRTree (*13*) to fulfill this task. It projects data points in a high-dimensional space to latent points in the low-dimensional space that directly form a tree structure. At the same time, the tree structure can incorporate the discriminative clustering information so that similar points are grouped together.

Let
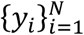
be a set of the *D*-dimensional column vectors consisting of ECI values corresponding to encounters of patients, where *N* is the total number of encounters. We define the latent points as
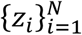
such that *z_i_* is the *d*-dimensional latent counterpart of *y_i_*. To transform *z_i_* to *y_i_*, a linear projection matrix *W* ( ℝ^*D*×*d*^ is used with *d* ≤ *D*. Moreover, in order to model the discriminative information via clustering, a set of latent points
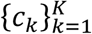
are introduced as the centroids of the latent points
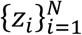
where *c_k_* ( ℝ^*d*^. The tree structure is represented by a connectivity indicator matrix *S* between cluster centroids with the (*k*, *k*′)th element as *s_kk′_*, that is, *s*_*k*,*k*′_ = 1 means that the *k*cth centroid and the *k*′th centroid are connected in the graph, and 0 otherwise. Specifically, a set of tree structures are denoted by *𝒮_T_*. By combining the above ingredients, DDRTree is formulated as the following optimization problem

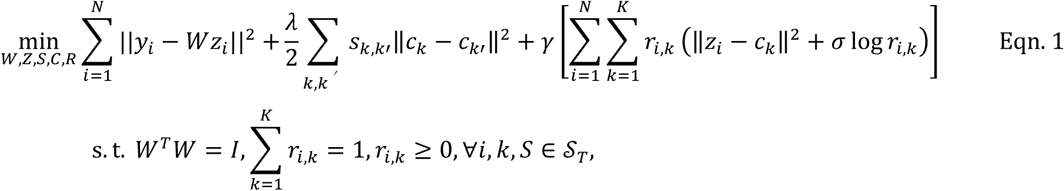

where the first term projects data into a low-dimensional space, the second term is responsible for structure learning, and the third term is interpreted as the objective function of soft K-means (*20*). Three hyper-parameters are added to connect the three terms: *λ* ≥ 0 is used for the reversed graph embedding, *γ* ≥ 0 is to balance the contribution of deterministic clustering, and *σ* ≥ 0 is to regulate the negative entropy regularization. Matrix *R* ( ℝ^*N*×*K*^with the (*i*, *k*)th entry as *r*_*i*,*k*_ can be interpreted as the probability of assigning *z_i_* to cluster *c_k_*. Details can be found in Supplement Method DDRTree. Details can be found in Supplement Methods (Section: DDRTree).

Due to the heterogeneity of data availability, we used a sub-group of patients with high data availability (the Learning cohort, n = 435) to understand disease progression trajectories and we projected data from the remaining patients (the Annotation Cohort, n = 1,146) to the learned trajectories for annotation and association analysis. The inclusion criteria of the Learning cohort were: ⩾ 50 encounters and every ECI had values from at least one encounter. Details can be found in Supplement Methods (Section: Imputation).

### Annotation of discovered clusters of clinical encounters

We comprehensively annotated each discovered cluster with corresponding clinical conditions including age, kidney function, *APOL1* genotype, hypertension, glucose control, obesity, liver function, and cardiovascular risk. Details are listed in Supplement Methods (Section: Annotation). Specifically, the enrichment of risk genotypes in discovered clusters was measured by robust z-test of the log fold changes of odds ratios against randomized controls (repeats = 10,000). In robust z-test, mean and standard deviation were substituted by median and median absolute deviation, respectively, to reduce sensitivity to outliers (*21*). The enrichment results were listed in Supplement Table S4.

### Classifying new encounters

To project each new patient’s encounters onto the learned structure and clusters, we still assume that the projection matrix *W*, the centroids *C*, and the graph structure *S* are the same as the learned variables from training data by solving the problem in Eqn. 1. Hence, the reformulated optimization problem for a new set of encounters
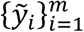
and the embedding points denoted by
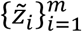
is formulated as

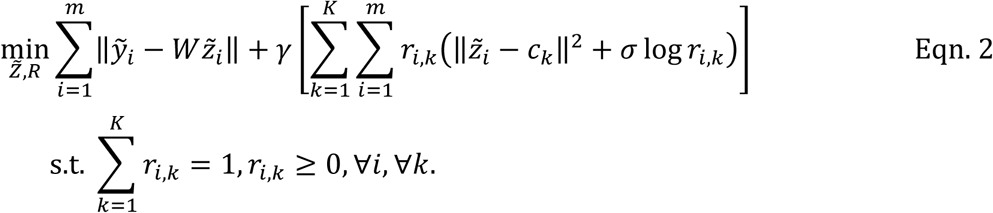

The objective function is

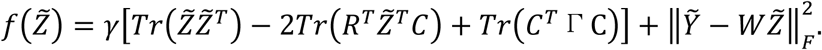

where Γ is a diagonal matrix with the (*k*, *k*)th entry as
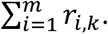
Problem in Eqn. 2 has the closed form solution by setting the first derivative to zero, we get

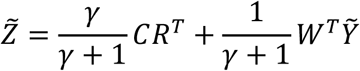

As a result, the alternating optimization algorithm can be used to solve this problem. Once the optimal *Z* and *R* are obtained, we can consider *Z̃_i_* as the low-dimensional projection of new patent *i* and *r*_*i*, *k*_ is the probability of assigning the *i*th encounter to cluster *k*.

The EMR data for the clinical encounters of Annotation Cohort (n = 1,146) was projected onto the learned progression trajectories and classified to the learned clusters.

### Prediction capability

To evaluate the power of predicting disease progression using Essential Clinical Indices EMR data and the learned disease progression trajectories, we used the k-nearest neighbors approach for prediction and leave-one-out for validation. The rationale was that patients with similar progression trajectories in the past were more likely to share similar future progression. For a given new patient with a sequence of encounters, we aimed to select a set of similar patients from the learning cohort. We leverage the progression tree structure learned by DDRTree model as a unified reference to the aligned sequence of the states of each patient. We developed a novel structure-based similarity function to measure the similarity between two patients. The basic idea was illustrated in Supplement Figure S1. For the obtained graph structure *S* and the clusters {*g*_1_,⋯,*g_r_*} from a set of visits of many patients, each cluster was denoted by the disease state and there were *r* states in total. Let *c*_*g*_1__, ⋯, *c_g_r__* be the cluster centers. Note that disease states are different from centroids *C* ∈ ℝ^*d*×*k*^ used in the DDRTree. The relationships among {*g*_1_, ⋯ ,*g_r_*}, *C* and visits of patients *X* were: each visit was assigned by one disease state, and each visit can also be mapped to its closest centroid on the tree. These relationships were useful to compute the evolutionary path via the tree structure given a sequence of visits of one new patient.

To accomplish this, we built a disease state tree via tree simplification from the 800 landmarks to 30 disease states (clusters) as shown in Supplement Figure S1 B, then aligned the clinical encounters of a patient to this tree to infer a sequence of disease states as shown in Supplement Figure S1 A. We defined the similarity between the disease state sequences of two patients according to the longest common subsequence. Thus, the simplified disease state tree could be used to predict disease progression using k-nearest neighbor approach.

We then performed an approximate leave-one-out validation to evaluate the performance of the learned trajectories in predicting disease progression. We took one patient out of the learning set as the query patient and the rest of patients as the database. For each query patient, we temporarily split the sequence of *t* visits ordered by the time into two sub-sequence at a split ratio of *r* ∈ [0,1]: the first subsequence with *r* ⋅ *t* disease states was used to predict the disease progression, and the latter subsequence with (1 – *r*) ⋅ *t* disease states was used to evaluate the prediction accuracy. This leave-one-out validation was approximate because the tree structure was not re-learned. We assumed that the overall tree structure would not change significantly when it was re-learned using the *n* – 1 patients from the learning cohort.

Mathematical details are provided in Supplement Material Methods (Section: Validation).

## Results and Discussion

### Demographical characteristics

The identification of the study cohort is outlined in Figure 1 and demographic characteristics are summarized in Table 1. As expected, patients in the Learning Cohort were followed longer (15.1 ± 4.77yr) than those in the Annotation Cohort (9.48 ± 6.48yr). Patients with high-risk *APOL1* genotypes were more likely to have more clinical visits and thus to be selected in the Learning Cohort, with percentages of 18.9% versus 14.3% if including patients with uncertain *APOL1* genotypes, and 21.0% versus 15.4% if only including patients with known *APOL1* genotypes. There were fewer males (43.6%) than females in our cohort, which was common for longitudinal studies of a midlife to senior age population (age: 57.2 ± 12.2yr). Interestingly, males were less represented in the Learning Cohort (39.5%) than the Annotation Cohort (45.1%) with a p-value of Chi square test ≤ 2.2e-16, suggesting that males had fewer clinic visits. Clinical conditions such as blood pressure and blood glucose level were similar in the two cohorts.

**Figure 1.**
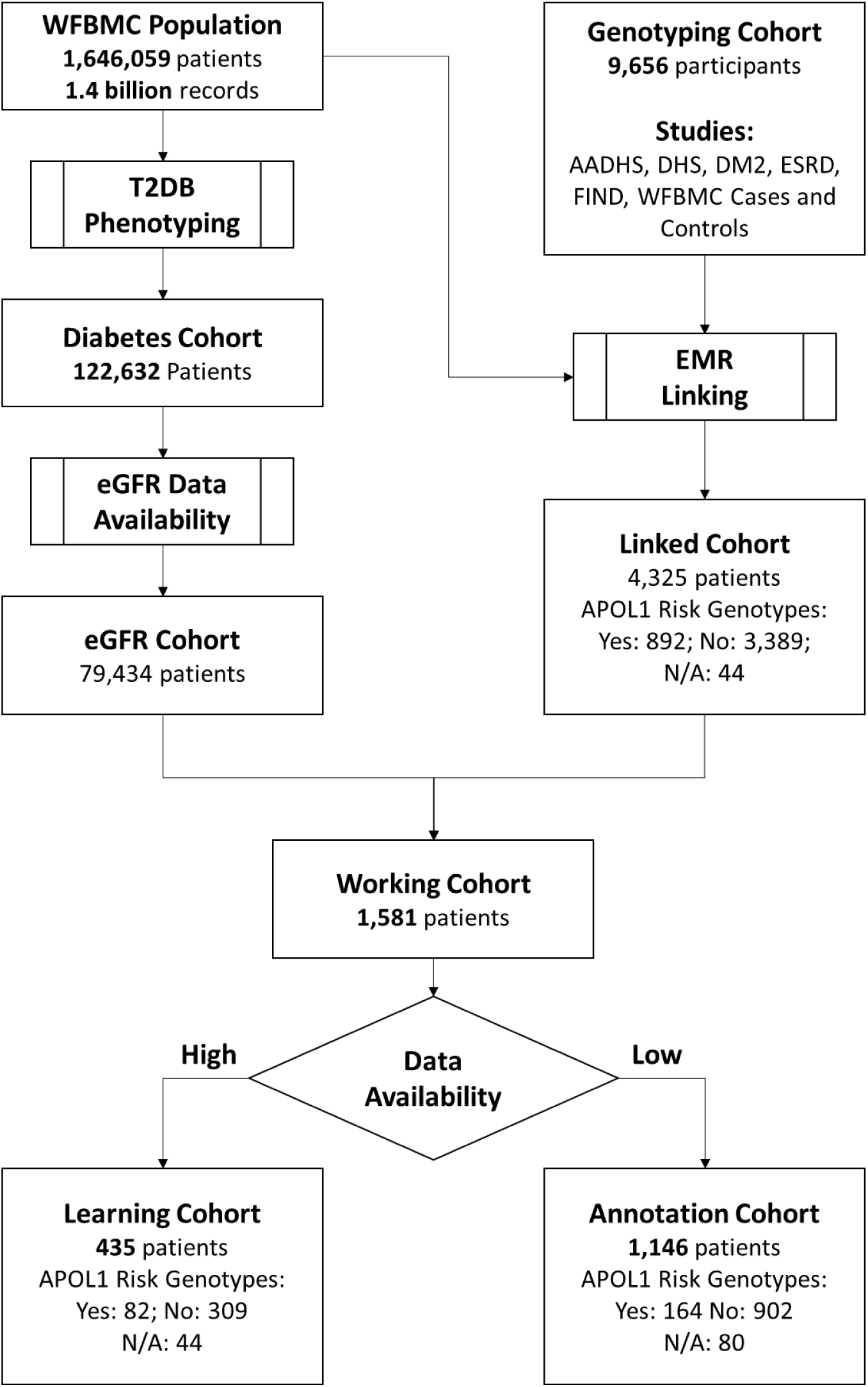
Outline of cohort identification.

**Table 1.**
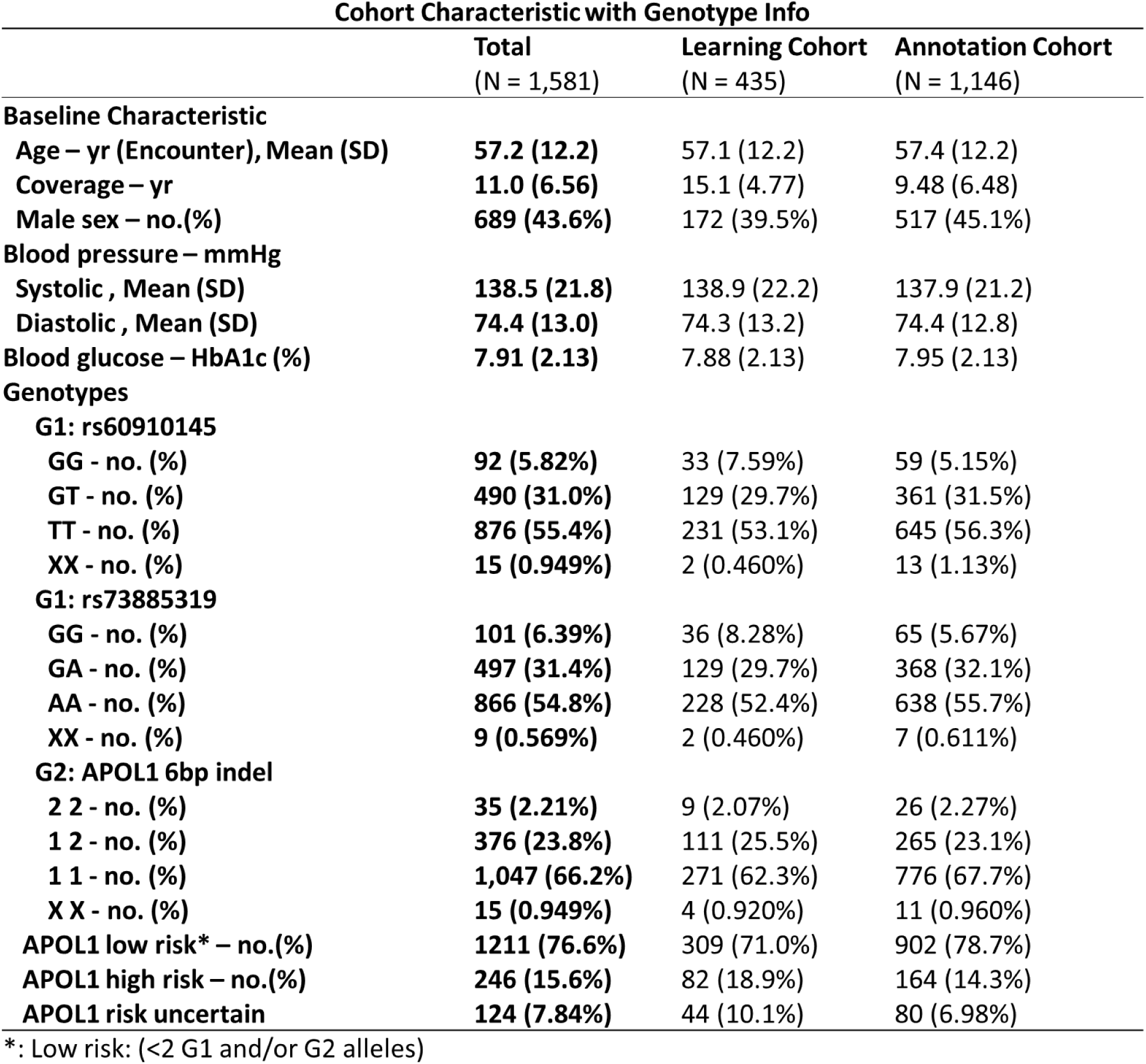
Cohort demographic characteristics.

### Data coverage

The EMR data in the 435 patients from the learning cohort was rich in records and features. They covered 58.75% of the overall encounters (49,357 of 84,009) for the working cohort of 1,581 patients and 58.11% (205,940 of 354,398) of Essential Clinical Index records. As shown in Figure 2, the learning cohort was intensively covered by EMR data. The longitudinal coverage (Figure 2 A) for each patient was 15.14 ± 4.77 years, with 13.79 ± 4.15 years having at least one clinic visit. The clinical encounters from the midlife-to-senior age ranges (35 – 80), which describe the development of a series of chronic diseases during aging are shown in Figure 2 B. Each patient had 113.46 ± 66.35 documented encounters (range 51 to 657) (see Figure 2 C and Supplement Figure S2). The most abundant Essential Clinical Indices were hemoglobin, TBIL, ALK, AST/SGOT, ALT/SGPT, blood pressure, eGFR, and weight, which were available in more than 10,000 encounters (Figure 2 D). Patients’ encounter counts and longitudinal coverage demonstrated weak correlation (Pearson’s product-moment correlation: 0.29, 95% CI: 0.23 to 0.36; p-value by t-test: 4.88e-15 Supplement Figure S3). That is, there were more visits the longer a patient stayed with our hospital for healthcare. Data completeness was low. As demonstrated in Figure 2 D and Figure 2 E, many of the Essential Clinical Indices were not available from most encounters. Data availability was 23.18% in the Learning Cohort and 4.27% in the Working Cohort. The availability of the Essential Clinical Indices showed strong correlations (Figure 2 E). The 18 clinical features formed four clusters, labeled C.1 through C.4.

**Figure 2.**
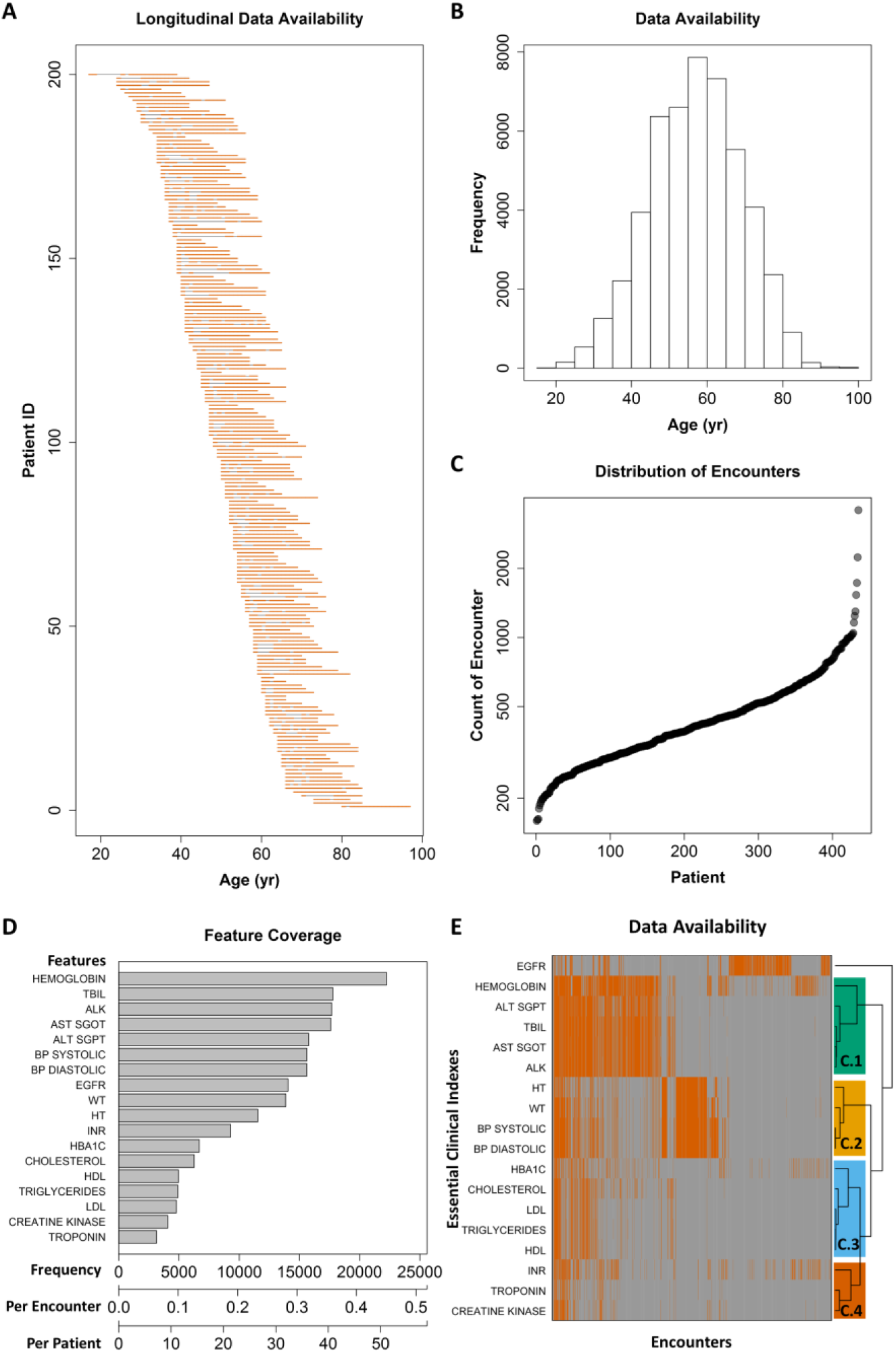
Data coverage of the Learning Cohort. (A) The distribution of longitudinal data availability for individual patients. Each line represent the encounter pattern of a patient, with orange sections for the years when a patient had EMR data, and gray section for years without visiting WFBH. Totally 200 randomly selected patients from the Learning Cohort were visualized for clarity. (B) Coverage of age range by EMR data at population level. (C) Distribution of encounters per patient. (D) The data availability of each Essential Clinical Indexes in terms of total counts, counts per encounter, and counts per patient. (E) The associations of data availability between Essential Clinical Indexes. Orange: available; gray: not available. Essential Clinical Indexes were clustered according to availability to four clusters, C.l through C.2.

For the overall cohort, EMR data was rich and abundant in longitudinal coverage and clinical features. The learning cohort intensively covered the trajectories of common clinical features over 45 years, from age 35 when early risk factors and symptoms of a chronic condition typically begins, up to age 80 when many chronic diseases have become end-stage. Data was abundant for recording the clinical health status accompanied with the progression of chronic diseases. A total of 49,357 encounters and 205,940 records were available for the 435-patient cohort, describing risks and status of chronic kidney disease, diabetes, hypertension, and cardiovascular disease. The rich EMR data provided deep learning of chronic disease progression from different perspectives and at multiple time scales. Meanwhile, EMR data were also sparse. The longitudinal coverage was nearly 14 years for most patients. Even for the most common clinical features (the 18 Essential Clinical Indices), the data availability was only 4.27%. This extreme sparseness challenges data analysis and machine learning approaches. To address these challenges, we used a subset of patients and encounters with more complete data to reliably understand disease progression patterns, and then mapped the remaining encounter data to the learned trajectories for further annotation. Meanwhile, the data availability showed strong patterns. The four clusters demonstrated in Figure 2 F represented different clinical practices. Essential Clinical Indices in cluster C.1 were often measured together for liver function. C.2 features were commonly measured at regular check-ups. Clinical assays in C.3 were often tested together to profile lipids and metabolic markers. Included in cluster C.4 were risk factors for cardiovascular disease.

### Chronic trajectories of health

The progression of chronic disease during aging was learned from the EMR-derived Essential Clinical Indices of 49,357 encounters on the 435-patient Learning Cohort using DDRTree (*13*). The trajectories (Figure 3 A) were represented in a latent graph space as a tree composed of 800 “landmarks” denoting the gradual development of clinical conditions. Encounters were clustered to 30 classes using k-means according to their locations on the trajectory tree (Figure 3 B). These encounter classes were annotated by patient age (Figure 3 C), kidney function (Figure 3 D), blood glucose control, hypertension, and BMI (Supplement Figures S4, S5, and S6, respectively).

**Figure 3.**
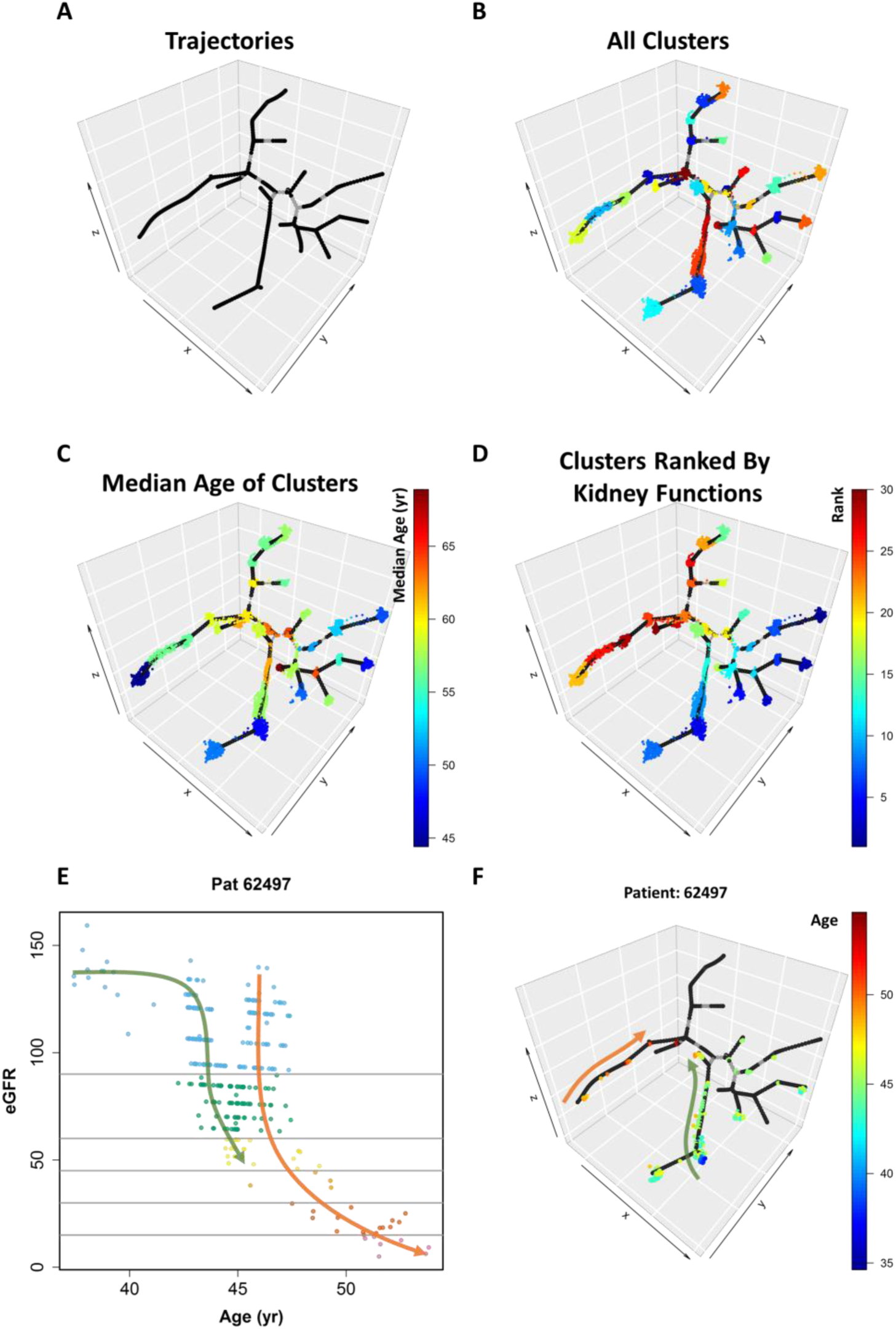
Chronic disease progression trajectories. (A) Learned trajectories represented by 800 landmarks. (B) Encounters (represented as dots) supporting the learned trajectories. Encounters were clustered into 30 classes and color coded. (C) Encounters color-coded with median age of corresponding clusters. (D) Encounters color-coded with ranked kidney functions of corresponding clusters, from normal kidney function (blue) to worst kidney function (red). (E) The progression of chronic kidney disease of a patient represented by estimated glomerular filtration rate (eGFR, unit: ml/min/1.73m^2^). (F) Encounter of the same patient mapped to the learned trajectories and color-coded according to age.

The disease progression in an individual patient could be visualized on the learned trajectory tree. As shown in Figure 3 E, a patient (Study ID: 62497) started to show kidney functional impairment at age 43, which quickly progressed to stage G3 at age 45 and recovered to normal at age 46. However, the patient subsequently developed rapid progression of kidney disease and advanced to kidney failure at age 52. Interestingly, when projected to the learned chronic trajectory tree (Figure 3 F), the progress of the CKD shifted from the low-risk trajectory (green trend line) to the high-risk trajectory (orange trend line) at age 46. Although kidney function apparently recovered at age 46, the nature of disease progression had changed.

The disease progression trajectories captured the gradual deterioration of kidney function. The directions of the progression of kidney disease along the trajectories (Figure 3 D) were consistent with aging (Figure 3 C). Multiple progression trajectories were discovered, reflecting the diversity of the disease. Meanwhile, when viewed from the perspective of individual patients, CKD progression could either remain on a major trajectory, or dynamically shift between trajectories, or switch trajectories. This exemplifies the complexity and heterogeneity of CKD development in populations. For patients who demonstrated competing trajectories and outcomes, further studies may reveal the risk factors that drove them toward trajectories of worse prognosis and suggest potential interventions to prevent changes in trajectory.

### Chronic trajectories in patients with *APOL1* renal-risk genotypes

Patients with *APOL1* high-risk genotypes were more likely to progress along a trajectory branch featured with classes 17, 10, and 18, with odds ratios higher than 2 (Figure 4 A). This branch was also associated with severe kidney function loss (Figure 4, Figure 3 D, Supplement Figure S7). In contrast, two low-risk branches (composed of classes 12, 7, 25 and 24, 16, 26, 9, respectively) were enriched with patients with non-renal-risk *APOL1* genotypes. During the clinical visits mapped to these two branches, the majority of patients demonstrated no sign of kidney disease (Normal or stage G1) or very early symptoms of kidney functional impairment (stage G2) (Figure 4, Figure 3 D, Supplement Figure S5).

**Figure 4.**
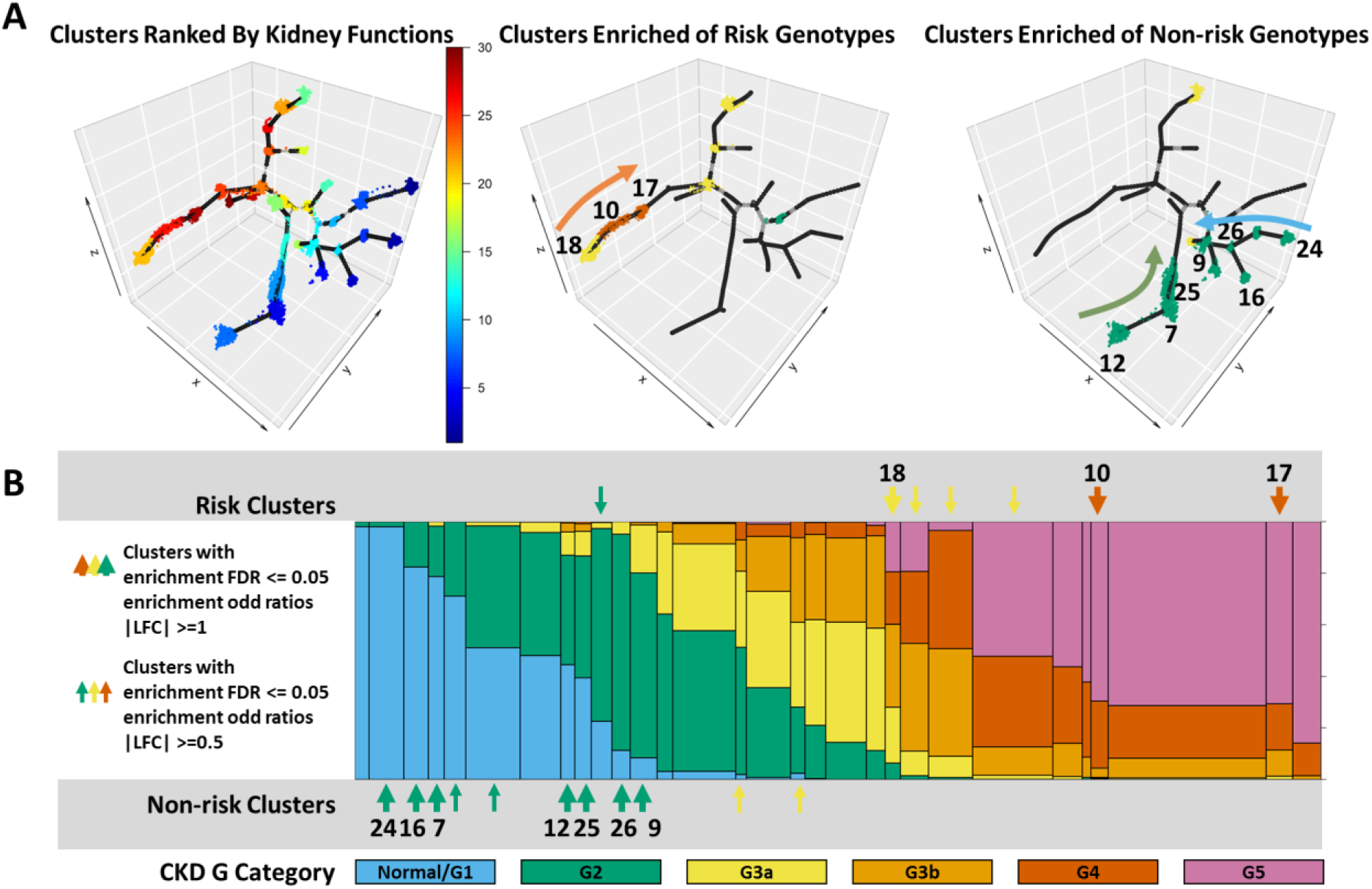
Enrichment of APOL1 risk genotypes on disease progression trajectories. (A) Left: the encounter clusters color-coded by kidney functions (blue: good, red: impaired); clusters enriched with encounters of patients of high- or low-risk APOL1 genotypes (middle and right, correspondingly). Encounter clusters of good, impaired, and severely damaged renal functions were colored coded in green, yellow, and red, respectively. The high-risk branch and the low-risk-1 and low-risk-2 branches were denoted by the orange, green, and blue arrows, respectively. (B) Encounter clusters were sorted by corresponding kidney functions, from good to impaired (middle). The distribution of chronic kidney disease stages in each encounter clusters were color-coded, from stage Gl/normal through G5. Encounter clusters enriched with high- or low-APOL1 risk genotypes were indicated by arrows.

We further examined whether the high-risk trajectory was associated with older age. Interestingly, the age distributions of these three branches were similar, with the high-risk branch slightly younger (age 51.95 ± 12.23, 52.82 ± 9.33, and 54.51 ± 12.45 for the high-risk and the two low-risk branches, respectively). This suggests that the more severe renal functional change in the high-risk trajectory is independent from age.

Acute kidney injury (AKI) is a known risk factor for CKD. We assessed the enrichment of AKI on the high-risk branch with respect to the two low-risk branches according to the ICD9 codes (584.^∗^). The odds ratios were 20.26 (95% CI: 15.01, 27.35) and 34.08 (95% CI: 25.49, 45.57), respectively (Supplement Table S5). However, the overall association between AKI and CKD was independent from *APOL1* renal-risk genotypes, with an odds ratio of 1.13 and p-value of 0.84 (Pearson’s Chi-squared test with Yates’ continuity correction). Stratified analysis (Supplement Table S5) further showed that patients without *APOL1* renal-risk phenotypes were more likely to change to the high-risk trajectory once they suffered an AKI event (odds ratios 61.5 and 49.2). This was consistent with the previously described case (Figure 3 E and F): patient 62497 had AKI at age 45.7 and at the same time changed from the low-risk to the high-risk trajectory. Patients progressing on the high-risk trajectory often experienced recurrent episodes of AKI. For example, patient 62497 experience 8 episodes of AKI at ages 47.5, 48.4, 48.8, 49.2, 49.5, 50.8, 51.4, and 52.6 before advancing to kidney failure. This is consistent with other studies on the concordance of AKI with CKD (*22*-*26*).

The genotype enrichment analysis strongly supported the validity of this study. Genotyping information was not used in trajectory learning, but patients with different *APOL1* genotypes were still enriched in different trajectories. This suggested that trajectories learned from the 18 Essential Clinical Indexes were able to capture kidney disease progression from different causes.

### Prediction of chronic disease progression

The predictive power of the learned trajectories for the projection of patient’s chronic health status using the kNN approach was compared with random control or with all neighbors by two metrics: proportion of good prediction and the Kolmogorov-Smirnov test of the cumulative distribution functions. We initially examined the proportion of patients with better predicted disease progress than random controls. As shown in Figure 5 A, at *k* = 7, the kNN model generated more accurate predictions in 87% of patients. The average similarity peaked at *k* = 9. The cumulative distribution functions of progression prediction by the kNN prediction model with 9 nearest neighbors (*k* = 9) or with all neighbors were shown in Figure 5 B. The trajectory-based kNN model demonstrated better prediction with p-value = 0.01255.

**Figure 5.**
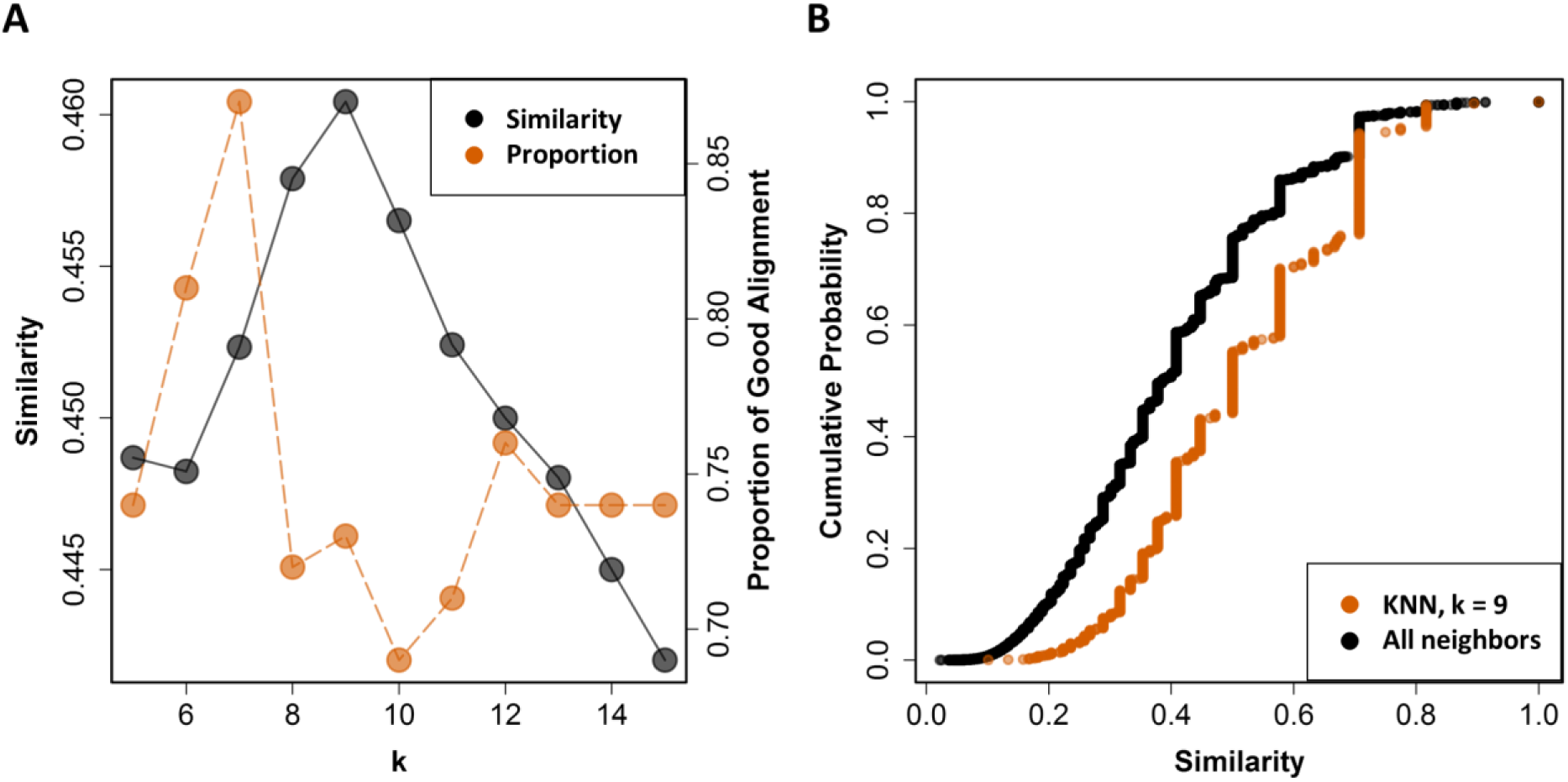
Prediction performance. (A) The average prediction similarity (gray) and the proportion of predictions with good alignment (orange) with respect to the number of nearest neighbors (*k*) were visualized. (B) The cumulative probability functions. Orange: for predictions using the k-nearest neighbor model (*k* = 9); black: using all samples.

The prediction power analysis was used to evaluate the potential of implementing the learned trajectories in projecting patient’s chronic health conditions. With a modest library size (n = 435), a simple trajectory-based similarity metric, and a non-parametric prediction model (kNN), the disease progression trajectories already demonstrated promising prediction power. The prediction power would be improved significantly with a larger library. Additional demographic and clinical features (*e*.*g*., age, sex, race, medication, diagnosis, and clinical procedures), widely available in the EMR, will significantly improve the prediction of patients’ health trajectories. Handling large libraries with multi-dimensional data are challenging. The performance of the kNN model also suggested that approximate nearest neighbor techniques such as k-d tree and locality sensitive hashing are promising for clinical use when cohort sizes are large. As suggested by Guo *et*. *al*., kNN predictive model shows good performance (power > 90% when effect size > 0.45) when the sample size of the corresponding class reaches 150 (*27*). That is, for any distinct trajectory pattern, when there are more than 150 patients in the cohort sharing this pattern, the prediction for new patients who also have this trajectory pattern will be accurate. Since patient numbers in EMR data can easily reach into the million range, it is promising to build a large database of trajectory patterns in existing patients and use locality sensitive hashing (*28*-*31*) and approximate nearest neighbor query techniques (*32*, *33*) to predict disease progressions in new patients.

## Acknowledgments

The authors acknowledge grants NIH R01 DK071891 (BIF), NIH R01 DK070941 (BIF), NIH R01 DK084149 (BIF), NIH U01 DK57298 (BIF), NIH R01 DK53591 (DWB), and National Science and Technology Support Program of China 2015BAK41B03 (XZ) for funding this study. The authors also acknowledge the DEMON high performance computing cluster, the Greengplum massively parallel processing database and the Data Lake cloud storage and computing facility at Wake Forest University School of Medicine, the Texas Advanced Computing Center (TACC) at The University of Texas at Austin (http://www.tacc.utexas.edu), and the Extreme Science and Engineering Discovery Environment (XSEDE, which is supported by National Science Foundation grant number ACI-1548562) for providing HPC resources that have contributed to the research results reported within this paper.

Author contributions JS, BIF, DWB, and MCYN initiated the study; JS and LW designed and performed the data analysis; XZ, LSH and JX prepared and preprocessed the data; FCH and SHC supervised the statistical analysis; JS and LW prepared the manuscript; BIF and DWB revised the manuscript.

